# Evidence of horizontal gene transfer and environmental selection impacting antibiotic resistance evolution in soil-dwelling *Listeria*

**DOI:** 10.1101/2024.06.27.600992

**Authors:** Ying-Xian Goh, Sai Manohar Balu Anupoju, Anthony Nguyen, Hailong Zhang, Monica Ponder, Leigh-Anne Krometis, Amy Pruden, Jingqiu Liao

## Abstract

Soil has been identified as an important reservoir of antibiotic resistance genes (ARGs) and there is a need to understand how corresponding environmental changes influence their emergence, evolution, and spread. As a soil-dwelling bacterial genus containing important pathogens, *Listeria,* including *L. monocytogenes*, the causative agent of listeriosis in humans, could serve as a key model for establishing this understanding. Notably, acquired antibiotic resistance among *L. monocytogenes* isolated from foods and the environment has been observed in some regions over the past decade. Here we characterized ARGs using 594 genomes representing 19 *Listeria* species that we previously isolated from soils across the United States. Among the five putatively functional ARGs identified, *lin*, which confers resistance to lincomycin, was the most prevalent, followed by *mprF, sul, fosX*, and *norB*. ARGs were found to be predominant in *Listeria sensu stricto* species and species more closely related to *L. monocytogenes* tended to harbor more ARGs. Notably, *lin, fosX,* and *norB* showed evidence of recent horizontal gene transfer (HGT) across species, likely through transformation as opposed to conjugation and transduction, while *mprF* and *sul* appear to have undergone positive selection. In addition, soil properties and surrounding land use were identified as the most important factors associated with ARG richness and genetic divergence, respectively. Using machine learning, we demonstrated that the presence of ARGs can be predicted from environmental variables with good accuracy (mean auROC of 0.76). Collectively, our data suggest that recent HGT and environmental selection played a vital role in the acquisition and diversification of ARGs in the soil environment.

## Introduction

Soil is a natural reservoir of antibiotic-resistance genes (ARGs), including ARGs that have been encountered in human pathogens^1, 2^, playing a pivotal role in the emergence, evolution, and dissemination of microbial antibiotic resistance across diverse ecosystems^3, 4^. *Listeria* is a soil-dwelling genus of bacteria^5^ that includes pathogenic members, such as *L. monocytogenes* and *L. ivanovii*. *L. monocytogenes* causes listeriosis in vulnerable human populations with a notable fatality rate of 20-30%^6, 7^, while *L. ivanovii* rarely causes listeriosis in humans and is primarily a pathogen of ruminant animals^8^. *Listeria* can be broadly divided into two groups: *sensu stricto* and *sensu lato*, based on the relatedness of species to *L. monocytogenes*, with *sensu stricto* species being more closely related^9^. The standard treatment for listeriosis is a combination of penicillin and aminopenicillins (ampicillin or amoxicillin)^9^ or ampicillin and gentamicin^10^. While the incidence of resistance among clinical *L. monocytogenes* to these antibiotics, fortunately, remains low at present, intrinsic resistance to cephalosporins exists, and increased resistance to penicillin, trimethoprim, and rifampicin has been observed^9, 11–14^. Furthermore, high rates of antibiotic resistance in *L. monocytogenes* have been observed in food-related environments^15^, possibly due to prolonged exposure to sublethal concentrations of antimicrobial agents in food processing and agriculture settings^16, 17^, such as poultries^15, 18^ and fresh produce factories^19, 20^. Since *Listeria* can be transmitted from soils directly to humans,^21^ or indirectly, via the food production chain^22^, they could be a key model for understanding how ARGs carried by soil microbes can be potentiated into human pathogens. Establishing a fundamental understanding of the ecological and evolutionary drivers of antibiotic resistance among soil-dwelling *Listeria* could help to better interpret current and future trends in antibiotic resistance patterns observed in clinical and food isolates. However, most studies of ARGs in *Listeria* have primarily focused on food-related and clinical isolates of *L. monocytogenes*^18, 23–25^, resulting in an incomplete understanding of the dynamics of ARGs in *Listeria* in the natural environment.

Previous studies indicate that environmental factors, including nutrient availability, temperature, pH, and exposure to natural or anthropogenic chemicals, can exert selective pressure favoring antibiotic resistance^26, 27^. Genes essential for metabolism and behavior, including ARGs, have been observed to undergo positive selection (PS) to adapt to varying environments^28, 29^. Apart from PS, environmental pressures can expedite horizontal gene transfer (HGT), a pivotal pathway for the evolution of new resistant strains^30^. HGT typically occurs through three mechanisms, transformation (i.e., the uptake of free DNA from the environment), conjugation (i.e., the direct transfer of genetic material from one bacterium to another through physical contact typically encoded by plasmids or transposons), and transduction (i.e., transfer of genetic material via viruses, such as prophages)^31^. Existing evidence suggests that the acquisition of ARGs among *L. monocytogenes* is mediated by HGT. For example, the acquisition of tetracycline and trimethoprim resistances in *L. monocytogenes* has been experimentally linked to transposons like Tn916-Tn1545 and Tn6198^32, 33^. Furthermore, a multi-drug resistant plasmid in *L. monocytogenes* can be naturally acquired by other closely related bacteria through transformation, thereby conferring resistance^34^. Despite the important role of the environment and HGT on antibiotic resistance, the prevalence of ARGs, the extent of HGT and PS acting on them, and the influence of environmental factors on the distribution and evolution of ARGs in soil-dwelling *Listeria* species remains largely unknown.

To bridge these knowledge gaps, we comprehensively examined the distribution of ARGs and associated HGT, PS, and environmental factors using in-depth population genetics analyses in a unique nationwide whole-genome dataset for 594 soil-dwelling *Listeria* strains representing 19 species, including *L. monocytogenes*. We identified five putatively functional ARGs (i.e., with coverage of > 80% and no premature stop codon detected), *lin*, *mprF, sul, fosX*, and *norB*, predominantly found among *Listeria sensu stricto* species. HGT of ARGs across *Listeria* species appeared to be mediated by transformation, rather than conjugation and transduction. We also revealed evidence of environmental selection acting on the richness and genetic divergence of ARGs, with machine learning models applied to predict the carriage of ARGs from environmental variables. This study yields new insights into the dynamics of antibiotic resistance in soils and suggests that environmental disturbance may facilitate the emergence and spread of ARGs among *Listeria* species.

## Results

### Prevalence and spatial distribution of ARGs among soil-dwelling *Listeria*

We identified seven distinct ARGs in soil-dwelling *Listeria*: *lin, mprF, sul, fosX, norB*, *dfrD,* and *mphB*, with the first five being functional and the latter two being non-functional (**Fig. 1a**). Specifically, *lin* confers resistance to lincomycin, *mprF* to defensin, daptomycin, and gallidermin, *sul* to sulfamethoxazole, *fosX* to fosfomycin*, norB* to fluoroquinolones and nalidixic acid*, dfrD* to trimethoprim, and *mphB* to erythromycin, telithromycin, quinupristin, pristinamycin IA, and virginiamycin S^35–37^. Among the functional ARGs, *lin* was most prevalent among *Listeria* isolates (82.66%) followed by *mprF* (82.32%), *sul* (81.14%), *fosX* (60.77%), and *norB* (58.42%) (**Fig. 1a**).

**Fig. 1.**
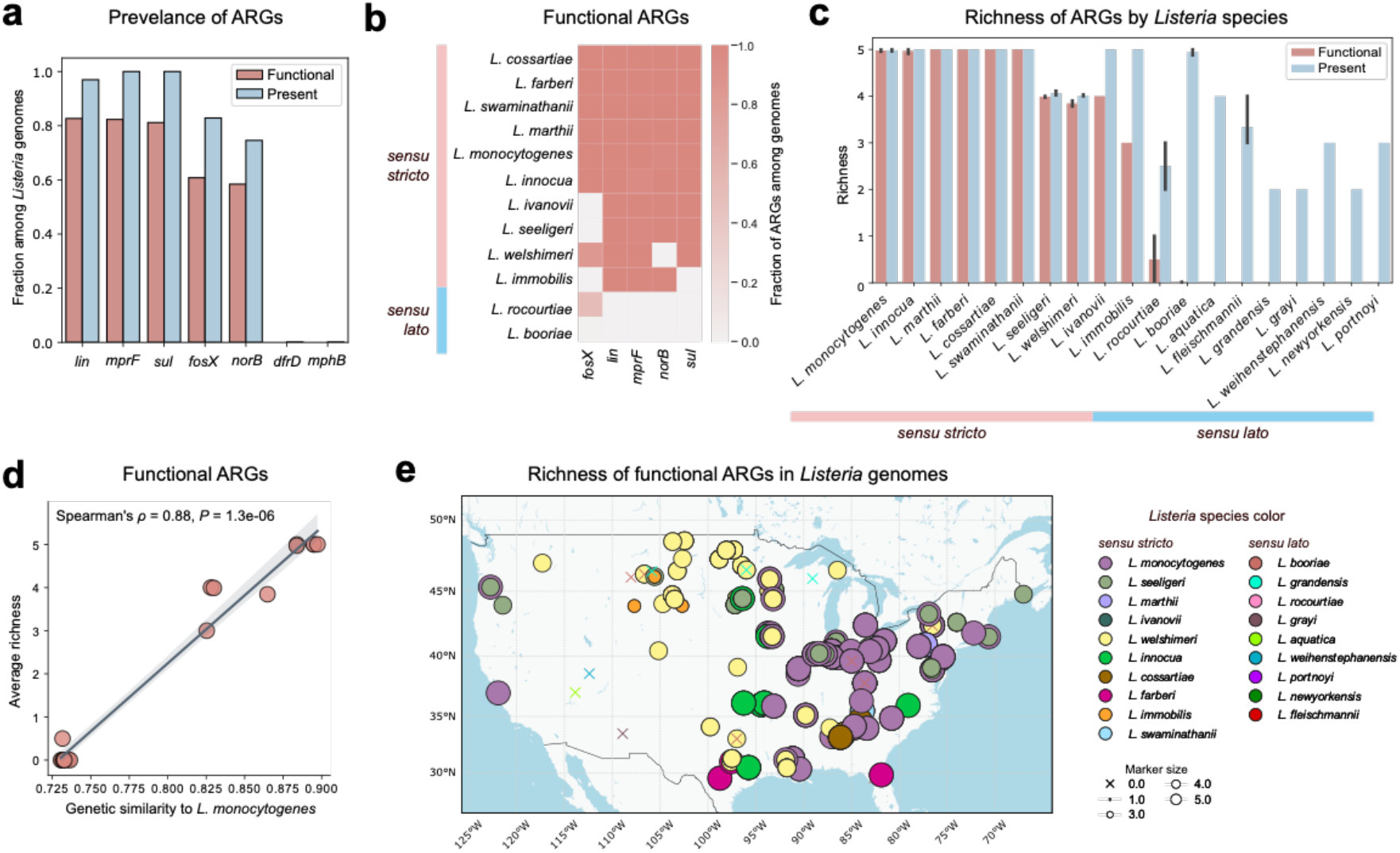
| ARG profiles among soil-dwelling *Listeria* and their national distribution. a Prevalence of both present (blue) and func4onal (red) ARGs across genomes. **b** Propor4on of func4onal ARGs among different *Listeria* species. **c** Richness of both present (blue) and func4onal (red) ARGs in different *Listeria* species. **d** Correla4on between gene4c similarity to *L. monocytogenes* and average richness of func4onal ARGs. Gene4c similarity was calculated based on pairwise ANI between different *Listeria* species and *L. monocytogenes*. ρ represents the Spearman’s rank correla4on coefficient. **e** Richness of func4onal ARGs among *Listeria* genomes across the United States. Circles and crosses indicate genomes with and without ARGs, respec4vely, and are color- coded by species. Circle size is propor4onal to the ARG richness calculated in each genome.

Overall, high richness of functional ARGs was consistently observed in all *sensu stricto* species, especially *L. monocytogenes, L. innocua, L. marthii, L. farberi, L. cossartiae*, and *L. swaminathanii*, but not in *sensu lato* species (**Fig. 1b**). While ARGs were present in *sensu lato* species, nearly all of them were non-functional and the overall prevalence was lower than *sensu stricto* species (**Fig. 1c**). Specifically, *lin, mprF,* and *sul* were consistently present in all strains of *sensu stricto* species (*n* = 491, **Supplementary** Fig. 1a), with each being functional in 100.00%, 99.59%, and 98.17% of these strains, respectively (**Fig. 1b**). Functional *fosX* was found in all *sensu stricto* species, except for *L. seeligeri, L. immobilis,* and *L. ivanovii.* Functional *norB* was found in all *sensu stricto* species, except for *L. welshimeri* (**Fig. 1b**). Among *sensu lato* species, functional ARGs were only detected in *fosX* in one *L. booriae* strain and one *L. rocourtiae* strain (**Fig. 1b**). Notably, ARG richness in *Listeria* species was highly correlated with their genetic similarity to *L. monocytogenes* for both present (Spearman correlation ρ = 0.88, *P* = 1.2e-06) (**Supplementary** Fig. 1b) and functional ARGs (Spearman correlation ρ = 0.88, *P* = 1.3e-06) (**Fig. 1d**). This indicates that species more closely related to *L. monocytogenes* tend to manifest a higher richness of ARGs.

Subsequently, we mapped the distribution of richness for both present ARGs (**Supplementary** Fig. 1c) and functional ARGs (**Fig. 1e**). Notably, eastern regions of the United States exhibited higher ARG richness compared to the western regions (**Fig. 1e, Supplementary** Fig. 1c). The geographic signal of ARG richness appeared to be driven by the distribution of species, given that *sensu stricto* species were more prevalent in the eastern regions, especially *L. monocytogenes,* which harbor a high richness of ARGs **(Fig. 1e, Supplementary** Fig. 1c).

### Evidence of HGT and PS of ARGs among soil-dwelling *Listeria*

To offer insights into the origin and HGT of the five functional ARGs (i.e., *lin, mprF, sul, fosX,* and *norB*), a gene tree was constructed for each ARG, depicted in **Figs. 2a-e**. If these genes were introduced through speciation, we anticipate a tree topology aligning with the phylogenetic relationships of *Listeria* species shown in **Fig. 3a**. However, if HGT occurred, the gene tree would not form distinct clusters based solely on species classification. Based on this assumption, the gene trees suggest that *lin* (**Fig. 2a**)*, fosX* (**Fig. 2b**), and *norB* (**Fig. 2c**) likely undergo HGT in certain species, particularly within the *sensu stricto* species, while *mprF* (**Fig. 2d**) and *sul* (**Fig. 2e**) may have arisen through the conventional process of speciation. For example, for *lin* (**Fig. 2a**), we observed clades that include a mix of *L. marthii* and *L. cossartiae* strains, as well as *L. innocua* and *L. farberi* strains. Similarly, *fosX* (**Fig. 2b**) displayed a comparable pattern among *sensu stricto* species, with clades containing a mix of *L. marthii* and *L. cossartiae* strains, as well as *L. welshimeri* and *L. monocytogenes* strains. Notably, the apparent HGT of *fosX* between *L. welshimeri* and *L. monocytogenes* strains serves as an example of gene transfer between pathogenic (*L. monocytogenes*) and non-pathogenic (*L. welshimeri*) species. Regarding *norB* (**Fig. 2c**), evidence of HGT was observed between *L. innocua* and *L. farberi* strains, as well as among *L. monocytogenes* (pathogenic), *L. innocua* (non-pathogenic), and *L. seeligeri* (non-pathogenic) strains.

**Fig. 2.**
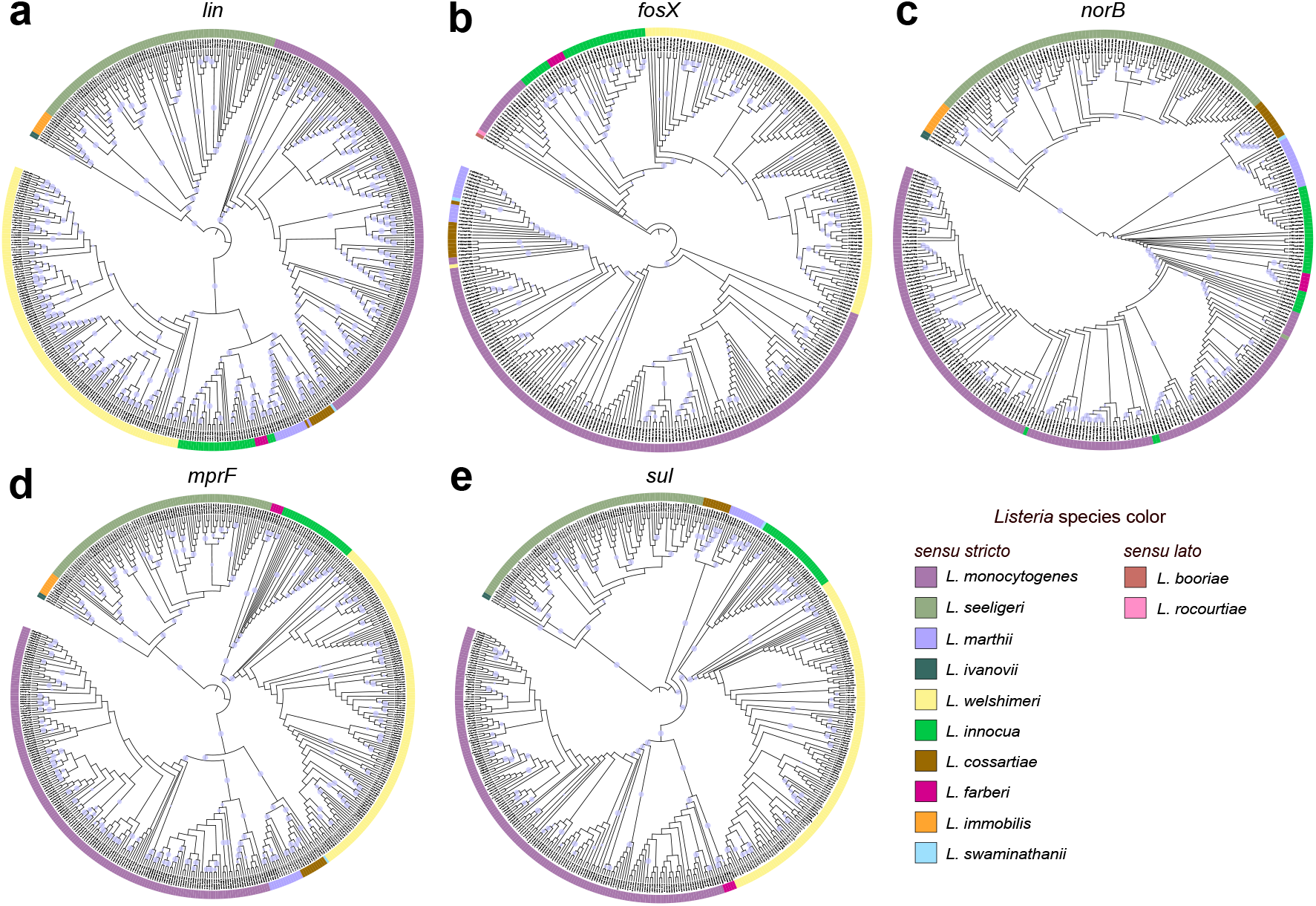
**| Maximum likelihood gene tree for functional ARGs detected among *Listeria* isolates.** a-e depicts the gene trees for **a** *lin*, **b** *fosX*, **c** *norB*, **d** *mprF*, and **e** *sul*, respec4vely, where evidence of HGT was observed in **a** *lin*, **b** *fosX*, and **c** *norB*, while PS is observed in **d** *mprF*, and **e** *sul*. The trees were constructed using sequences for *lin*, *fosX*, *norB*, *mprF*, and *sul* detected in 491, 361, 347, 489, and 482 genomes, respec4vely, with 1,000 bootstraps. The evolu4onary models used for construc4ng the tree were TN+F+I+R4, HKY+F+I+R3, TVM+F+I+R4, GTR+F+I+R5, and K3Pu+F+I+R3 for *lin, fosX, norB, mprF,* and *sul*, respec4vely. The trees were rooted by the midpoint. Bootstrap values >80% are indicated by light blue circles.

**Fig. 3.**
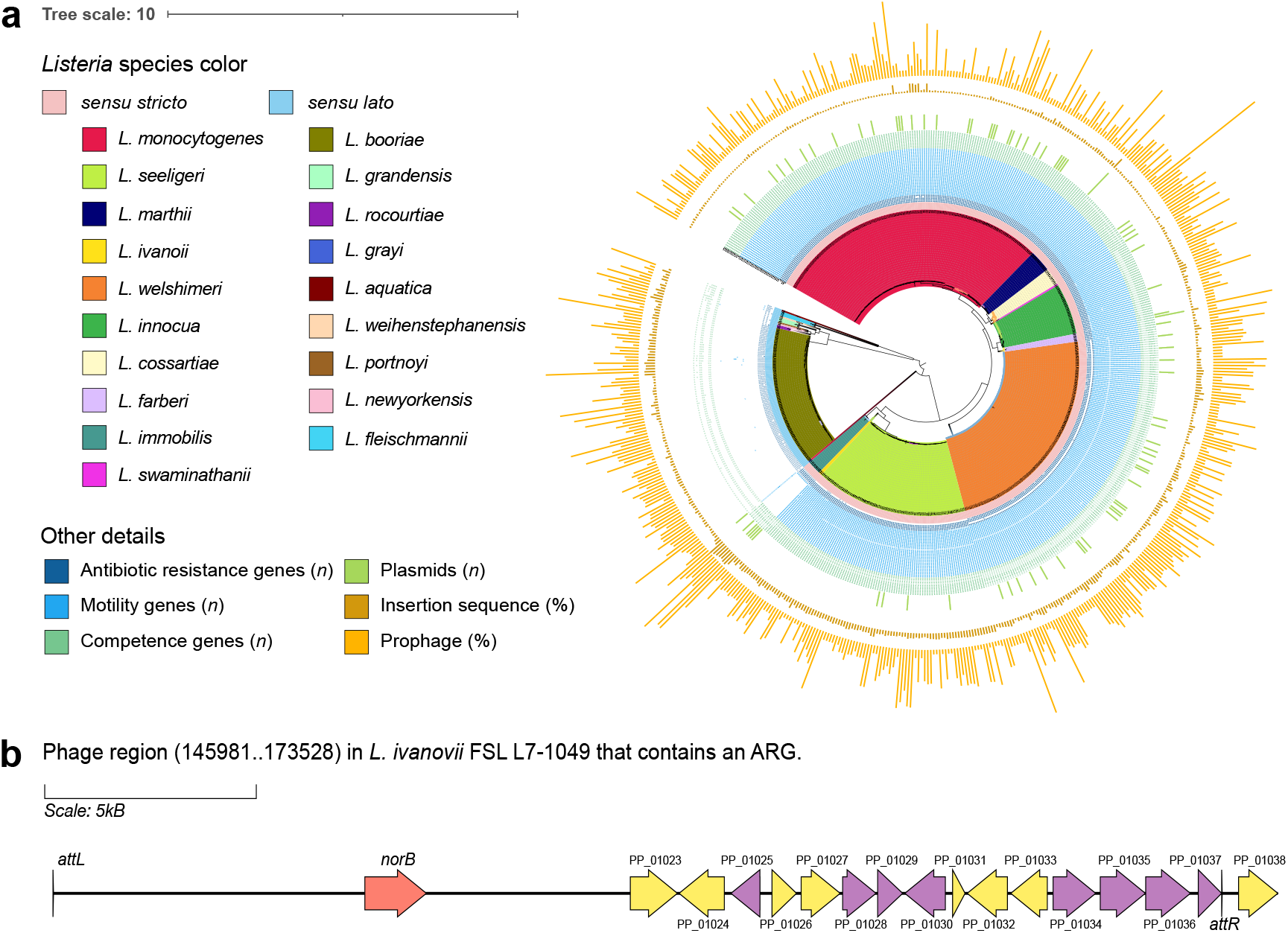
**| Overview of *Listeria* species phylogenies, motility genes, competence genes, and MGEs**. **a** The maximum likelihood phylogenetic tree was adapted from a previously published phylogenetic tree for *Listeria*, constructed from 594 genomes using core SNPs with 200 bootstraps^29^. The tree, rooted by the midpoint, features branches color-coded by *Listeria* species. Annotations include the status (presence/absence) of motility and competence genes, as well as MGEs (plasmids, ISs, and prophages). In the annotations, a filled box indicates the presence of a functional gene, an empty box indicates a non-functional gene (i.e., truncated or with premature stop codons), and a white box indicates the absence of the gene. **b** Visualization of a prophage carrying a *norB* gene in *L. ivanovii*. *norB*, genes encoding phage-related proteins, and genes encoding hypothetical proteins are shown in red, yellow, and purple, respec4vely. The lee and right afachment sites for the phage are referred to as *a8L* and *a8R*, respec4vely.

To understand if maintaining ARGs may offer advantages in fitness to adapt to certain environmental stressors, we performed a gene-wide test for PS for each of the five functional ARGs. Results showed that *mprF* (**Fig. 2d**, LRT = 13.932, *P* = 0.0004718) and *sul* (**Fig. 2e**, LRT =17.539, *P* = 7.77e-05) exhibited a better fit in the alternative unconstrained model (allowing for PS) compared to a null model. This suggests the presence of positive selection in *mprF* and *sul*, which were the only ARGs universally present among all 594 *Listeria* strains, i.e., evidence that these genes confer an advantage under environmental pressures. For the remaining ARGs – *lin*, *fosX*, and *norB* – there is no evidence supporting the presence of PS acting on these three genes (*P* > 0.05 for all).

### The probable mechanism of HGT of ARGs among soil-dwelling *Listeria*

To investigate potential mechanisms underlying the HGT in ARGs of *Listeria* in soil, we employed a predictive approach focusing on MGEs, including prophages, IS, composite transposons, and plasmids. The presence of ARGs within these MGEs could indicate their role as carriers, informing specific mechanisms of HGT. Among the 594 genomes examined, 1,338 prophages were identified (**Fig. 3a**). Out of these prophages, only 14.7% (*n* = 197) were classified as ‘intact’. Of note, we found 12 ‘incomplete’ prophages in *L. booriae*, 11 of which presented *lin* and one of which presented *norB* (**Supplementary Table 1**). However, these ARGs appeared to be not functional (**Fig. 3a**). The presence of remnants of ARGs in the prophages suggests a historical role of bacteriophages in transferring ARGs from other species to *L. booriae.* Apart from these observations, the only instance of recent HGT of a functional ARG potentially mediated by transduction was the *norB* gene located within a prophage, PHAGE_Bacill_SPbeta_NC_001884, of *L. ivanovii* L7-1049, a pathogenic species (**Fig. 3b, Supplementary Table 1**). The prophage region, delimited by the left (*attL*) and right (*attR*) attachment sites, comprises the *norB* gene and genes encoding other phage-related proteins and hypothetical proteins (**Fig. 3b**). The *norB* gene identified in this prophage was positioned in the first split branch in the *norB* tree (**Fig. 2c**), adjacent to the *norB* of another *L. ivanovii* strain L7-0121 rather than strains of a different species. This proximity suggests the occurrence of transduction events within this species, not across species.

A total of 4,023 ISs were identified (**Fig. 3a**), with only 18.3% (*n* = 735) being classified as ’complete’. IS3 and IS1182 families constituted 66.4% (*n* = 488) and 15.5% (*n* = 114) of the complete IS, respectively. Using ISAbR_finder, we identified an IS-associated ARG that matched with the functional *fosX* located on the negative strand in *L. welshimeri* L7- 1846, showing 100% identity and coverage. The copy of the IS3 transposase was located downstream of *fosX* on the positive strand, but IS3 transposition involves a copy-out-paste-in process that requires at least two copies of IS3^38^. Thus, we expect that the IS3 transposase under the configuration that it was found would not be sufficient for gene transfer. In addition, no composite transposons carrying ARGs were detected. For plasmids, only 81 were identified across the 594 draft genomes (**Supplementary Table 2**). The most dominant plasmid incompatibility (Inc) family and group were Inc18 (98.8%, *n* = 80) and repUS25 (96.3%, *n* = 78), respectively. Still, none of the plasmids identified in this study were found to carry any ARGs.

Given that conjugation and transduction did not appear to be substantial contributors to the HGT of functional ARGs across *Listeria* species, we further explored whether natural transformation might play a role. We predicted the presence of competence genes, which signify a bacterium’s capacity to uptake foreign DNA, or extracellular DNA (eDNA), from its surroundings and integrate it into its genome^39^. *Listeria sensu stricto* species generally possessed significantly more competence genes than *Listeria sensu lato* species (adjusted Mann-Whitney *U P* < 0.05 for all; **Fig. 3a, Supplementary Table 3**). Among *Listeria sensu stricto* species, competence genes were uniformly present, with over 90% functionality observed for several key competence genes, including *comK, coiA, cinA, comEC,* and *comEB* (**Fig. 3a**). On the contrary, among *Listeria sensu lato* species, specific competence genes such as *comGD, comG,* and *comGF* were completely absent (**Fig. 3a**). The sole functional competence gene among *Listeria sensu lato* species was *comEC*, identified in three *L. fleischmannii* strains (L7-1629, L7-1641, and L7-1645).

As motility plays a crucial role in enabling bacteria to move, providing adaptive advantages in new environments^40^ and facilitating DNA uptake^41^, we further predicted motility genes across the *Listeria* genomes. Consistent with competence genes, *Listeria sensu stricto* species possessed significantly more motility genes compared to *sensu lato* species (adjusted Mann-Whitney *U P* < 0.05 for all; **Fig. 3a, Supplementary Table 3**), which may increase their chance of being exposed to diverse DNA pools in the environment. Specifically, all 31 motility genes were present in every *Listeria sensu stricto* species, with 90% of these genes being identified as functional in more than 99% of strains, except for *L. immobilis*, the sole *sensu stricto* species identified thus far that lacks motility^42^. In contrast, almost none of the genomes among *sensu lato* species harbored any functional motility genes. Since functional ARGs and their HGT events were predominately observed among *Listeria sensu stricto* species, these collective observations regarding the distribution of competence and motility genes suggest that natural transformation may play a substantial role in the HGT of ARGs among *Listeria sensu stricto* species.

### Environmental factors associated with the richness and genetic divergence of ARGs among soil-dwelling *Listeria*

To assess the influence of the environment on ARG acquisition and evolution, we performed correlation analyses between environmental variables and ARG richness and sequence diversification. Considering the high correlation between ARG richness and genetic similarity of isolates to *L. monocytogenes* (Spearman ρ = 0.88, *P* < 0.001; **Fig. 1d**, **Supplementary** Fig. 1b) and a geographic signal likely driven by species (**Fig. 1e**, **Supplementary** Fig. 1c), genetic similarity could potentially confound identification of true correlations among environmental variables and ARG richness. Thus, Spearman partial correlation analysis, controlling for the genetic similarity of *Listeria* species to *L. monocytogenes,* was performed to investigate the relationship between ARG richness and 34 environmental variables. Thirteen environmental variables were significantly correlated with ARG richness (adjusted Spearman *P* < 0.05 for all). Among these, seven variables, including aluminum, forest, zinc, manganese, iron, longitude, and developed areas with < 20% impervious surface, exhibited a positive correlation, while the remaining six: copper, wetland, molybdenum, magnesium, pH, and potassium, showed a negative correlation (**Fig. 4a**). To further quantify the contributions of different environmental variable categories to the observed variation in ARG richness, VPA was conducted. To exclude potential confounding effects, genetic similarity to *L. monocytogenes* was also included in this analysis. Of note, the climate category was excluded because no climatic variables were found to be significant (**Fig. 4a**). As expected, genetic similarity alone accounted for the majority of the variation (adjusted *R*^2^, 80.53%) (**Fig. 4b**). Among the environmental variables, soil properties explained 7.37% of the variation, similar to the 6.74% explained by land use (**Fig. 4b**). Geolocation accounted for <1% of variation (**Fig. 4b).** MDS analysis further demonstrated that isolates with and without functional ARGs formed significantly different clusters based on environmental conditions (**Supplementary** Fig. 2; PERMANOVA *P* < 0.001). As functional ARGs were predominately detected in *Listeria sensu stricto* species (**Fig. 1c**), we hypothesized that environmental variables significantly differ between *Listeria sensu stricto* and *sensu lato* species. Indeed, latitude, minimum temperature, coverage of pasture, cropland, and barren areas were found to be significantly higher for *Listeria sensu stricto* species, while coverage of shrubland, maximum temperature, precipitation, and several soil properties (aluminum, copper, iron, molybdenum, and moisture) were found to be significantly higher for *Listeria sensu lato* species (adjusted Mann-Whitney *U P* < 0.05 for all; **Fig. 4c**). These results collectively suggest that environmental conditions, particularly soil properties, play a role in ARG acquisition in *Listeria*.

**Fig. 4.**
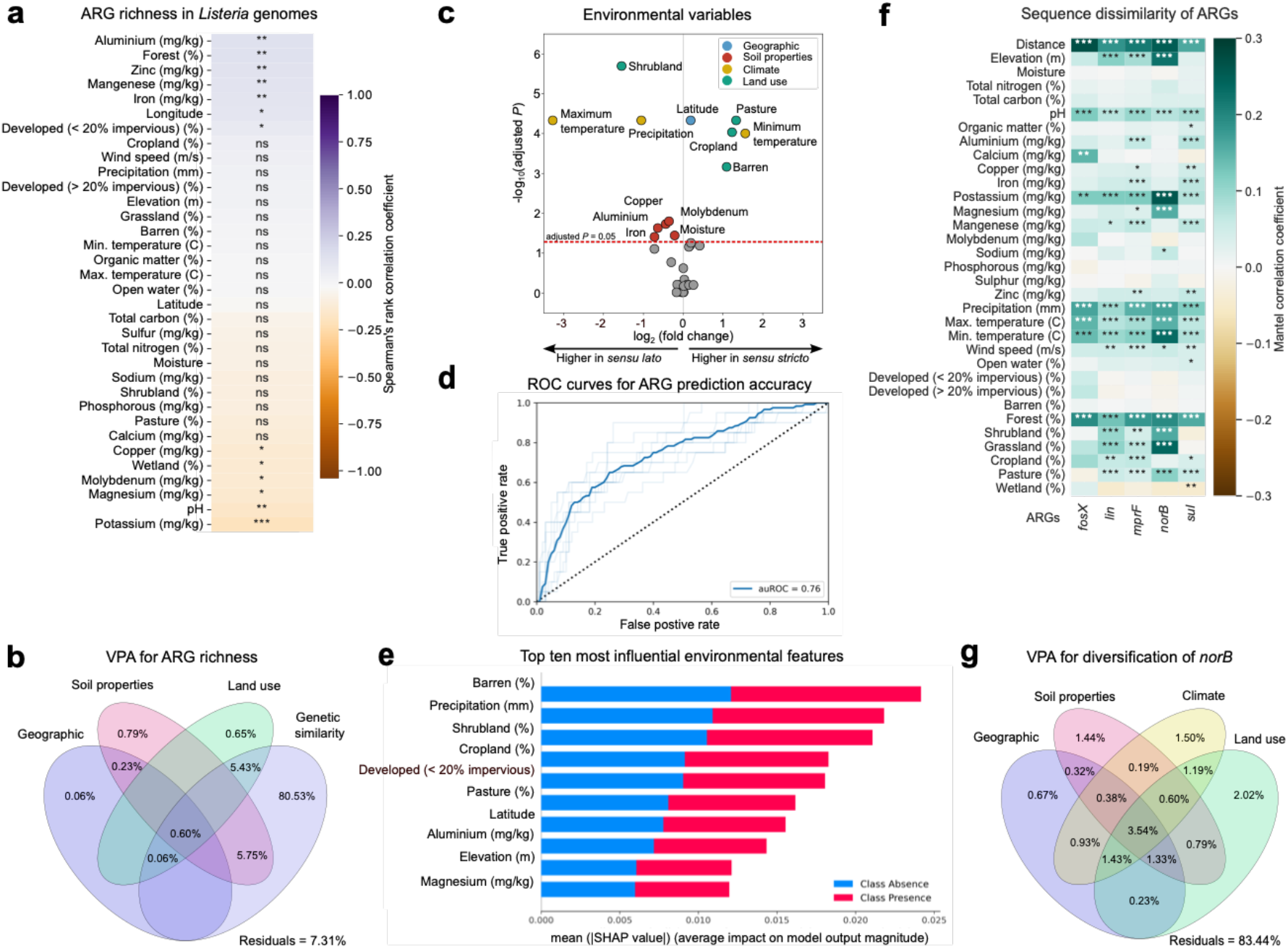
| Environmental conditions associated with ARG richness and genetic divergence among the *Listeria* isolates. a Spearman’s par4al correla4on between ARG richness in *Listeria* genomes and environmental variables, controlled for gene4c similarity for *L. monocytogenes*. **b** Venn diagram of varia4on par44oning analysis (VPA) showing the varia4on of the richness of func4onal ARG being explained by the environmental variable groups with significant correla4ons detected (i.e., geographic loca4ons, soil proper4es, and surrounding land use) and gene4c similarity for *L. monocytogenes*. Residuals indicate unexplained varia4on. **c** Volcano plot illustra4ng the significance (two-sided Mann-Whitney; y-axis) of the difference between environmental variables (fold change; x-axis) among *Listeria sensu stricto* and *sensu lato* species). Variables above the red dashed line have an adjusted *P* < 0.05. Dots are color-coded by environmental variable types. **d** ARG predic4on accuracy of random forest model based on environmental variables (mean auROC = 0.76). The light blue lines show ROC curves for models trained only on the training set. Each curve reflects one evalua4on using holdout data, repeated 10 4mes. The dark blue line represents the mean performance across these repe44ons. **e** The top ten most predic4ve features on overall predic4on of ARG presence/absence in **d** (SHAP-based; *X* axis), sorted by descending importance. **f** Mantel tests between ARG sequence dissimilarity and environmental variables. **g** Venn diagram of VPA showing the varia4on of the gene4c divergence of *norB* explained by environmental variable groups. For **a** and **f**, significance levels are denoted by “*”, “**”, “***”, and “****” for adjusted *P* < 0.05, *P* < 0.01, *P* < 0.001, and *P* < 0.0001, respec4vely, and “ns” indicates not significant.

Given the relationship between environmental variables and ARG richness that we observed, we hypothesize that the status (presence/absence) of ARGs in *Listeria* is predictable using environmental variables. To test our hypothesis, we compared different machine learning algorithms (see Methods) and identified random forest as the best algorithm for predicting the presence of ARGs with environmental variables as features. The model utilized logistic loss to assess the quality of splits, had a maximum depth of 3, considered the logarithm base 2 of the total number of features to determine the best split, and comprised 800 trees in the forest. After hyperparameter tuning (**Supplementary Table 4**), the best model achieved a mean auROC of 0.76 (**Fig. 4d**) and a mean auPR of 0.88 (**Supplementary** Fig. 3). To interpret outputs from the best machine learning model, we utilized SHAP^43^ to assess the importance of each feature to the prediction. The ten most influential environmental factors were barren land, precipitation, shrubland, cropland, developed areas with < 20% impervious surface, pasture, latitude, aluminum, elevation, and magnesium (**Fig. 4e**). These findings underscore the predictive capability of environmental factors for determining the presence of ARGs in *Listeria* species.

Lastly, to investigate the interplay between the genetic divergence of ARGs and environmental factors, Mantel tests were conducted to assess correlations between the sequence dissimilarity of each ARG and the distance of each environmental variable. We identified seven universal variables – geographic distance, pH, potassium, precipitation, maximum and minimum temperatures, and surrounding forest coverage – which exhibited a consistent significant positive correlation with sequence dissimilarity across all five functional ARGs (Mantel *P* < 0.05 for all; **Fig. 4f**). Among these ARGs, *mprF* exhibited correlations with the greatest number of significant environmental variables (*n* = 19). To delineate the contribution of environmental variable groups to the observed variation in ARG sequence dissimilarity, we further conducted VPA. Among the ARGs, *norB* (**Fig. 4g**) sequence divergence was the most affected by environmental variables, accounting for 16.56% of the explained variation. For *fosX*, *mprF*, *lin*, and *sul*, environmental factors contributed to 12.99%, 7.93%, 6.79%, and 6.26% of explained variation (**Supplementary** Figs. 4a-d), respectively. Despite the varying contributions of environmental variables to the sequence divergence of different ARGs, a consistent pattern emerged that surrounding land use was the most influential factor across all five ARGs, independently (and collectively) explaining 2.02% (11.13%), 3.04% (6.48%), 1.35% (4.77%), 1.57% (4.42%), and 1.48% (3.45%) of the variation for *norB, fosX, mprF, lin,* and *sul,* respectively, compared to other environmental variable groups. Collectively, these results suggest that, like richness, the genetic diversification of ARGs might be influenced by environmental conditions as well. However, instead of soil properties, surrounding land use tends to play a more important role in the diversification of ARGs.

## Discussion

### ARGs are predominantly present in *Listeria sensu stricto* species in soils, with limited evidence of the impact of antibiotics used in clinical settings

This study explored the dynamics of ARGs within soil-dwelling *Listeria* species, including *sensu stricto species*, which are more closely related to *L. monocytogenes*, and *sensu lato* species. We identified five functional ARGs, *lin*, *mprF*, *sul*, *fosX*, and *norB*, predominantly in *Listeria sensu stricto* species. Most of these five ARGs are still present in *Listeria sensu lato* species, but appear to be non-functional, suggesting that carrying ARGs may cause metabolic costs in these species^44^. In contrast, maintaining at least some of these ARGs in the genomes might increase fitness in *Listeria sensu stricto* species evidenced by the PS in two ARGs, *mprF* and *sul*, in these species. The large discrepancy in the prevalence of ARGs between *Listeria sensu stricto* and *sensu lato* species may also be partly attributed to the observed different conditions in the soil environment they encounter.

The five ARGs, *lin*, *mprF*, *sul*, *fosX*, and *norB*, identified in soil-dwelling *Listeria* species in this study are considered intrinsic because they are part of the core genome for *L. monocytogenes*^23, 35^. The current treatment protocol for listeriosis involves a combination of penicillin and aminopenicillins (ampicillin or amoxicillin)^9^ or ampicillin and gentamicin^10^. ARGs conferring resistance to these antibiotics used in clinical treatment, including *ampR*, *aacA4,* and *aadC*, were not detected in *Listeria* soil isolates included in this study, which suggests that in soil-dwelling *Listeria* examined in this study have not been influenced by antibiotics used in clinical settings. Indeed, findings from ARG surveillance of *L. monocytogenes* in food-related and clinical settings in France^23^, Denmark^45^, and Spain^46^ consistently report that acquired resistance is limited in *L. monocytogenes* and this pathogen remains susceptible to antibiotics over time. Therefore, while emerging resistance in *L. monocytogenes* is observed for certain clinical-use antibiotics, like penicillin^11^ and rifampicin^14^, resistance to the antibiotics used in patients with listeriosis (aminopenicillins and gentamicin) remains rare^9, 23^.

### ARGs in *Listeria sensu stricto* species show evidence of HGT, likely caused by natural transformation

HGT events were observed in *lin, fosX,* and *norB* among *Listeria sensu stricto* species, including between pathogenic (*L. monocytogenes*) and non-pathogenic species (*L. welshimeri, L. innocua*, and *L. seeligeri*) in this study. This observation supports the notion that HGT tends to display a bias toward individuals and species that are more closely related^47^. HGT of ARGs has also been observed among both clinical and food isolates from various *L. monocytogenes* clonal complexes, with tetracycline resistance identified as the most prevalent acquired resistance phenotype^48^. This has been primarily attributed to the presence of composite transposons like Tn916-Tn1545 carrying tetracycline resistance (Tn916-carrying *tetM* genes) in *L. monocytogenes*^23, 32, 33^. However, we did not identify any tetracycline resistance genes (*tetM* and *tetS*) among the *Listeria* soil isolates examined in this study. This is likely attributed to the widespread use of tetracycline in clinical and food-related environments^49–51^, whereas the baseline tetracycline concentrations in less disturbed environments might be low^52^.

Given the limited instances of acquired resistance observed from transposons, prophages, or plasmids in this study, we propose that transduction and conjugation may not be the primary mechanisms for the HGT of ARGs observed in *Listeria* soil isolates. Instead, we found that natural transformation is the most likely mechanism. Natural transformation relies on the recipient bacterium that expresses the competence machinery^53^ and largely depends on the uptake and incorporation of exogenous naked DNA from the environment into the genomes of competent recipient organisms^54^. We detected a complete set of competence genes that enable the acquisition of foreign DNA in most of the genomes of *Listeria sensu stricto* species, where HGT events of ARGs were observed. However, despite the presence of genes associated with competence machinery, *L. monocytogenes* is not recognized as naturally transformable in lab settings^55^. The absence of competence in *L. monocytogenes* has been attributed to the truncation of the *comK* gene, which cleaved into two parts by a 42-kb region containing several ORFs encoding phage-related products^56^. The regulation of the Com system relies on the formation of a functional *comK* gene via prophage excision^57^. In this study, based on the high coverage (> 80%) of *comK* genes detected in the genomes, this gene does not appear to be truncated in *Listeria sensu stricto* species. Nevertheless, even with strains containing an intact *comK* gene, attempts at transforming *L. monocytogenes* have been unsuccessful under lab conditions^56^. This suggests that *Listeria* may require unusual conditions (beyond competence minimal medium, at 37°C, and selection on BHI agar supplemented with chloramphenicol, the specifics of which are still unknown) for competence^56^, which complex soil environments may uniquely be able to provide, facilitating HGT of ARGs in *Listeria sensu stricto* species via natural transformation.

### Environmental selection plays a role in the acquisition and diversification of ARGs among soil-dwelling *Listeria*

Understanding the associations between environmental factors and ARGs is crucial for unraveling the dynamics and evolution of antibiotic resistance under the context of climate change. In this study, we observed that the richness of ARGs in soil-dwelling *Listeria* is predominantly associated with soil physicochemical properties, such as pH, aluminum, zinc, manganese, iron, copper, magnesium, and potassium. Consistent with this finding, previous studies reported that nanoalumina can enhance the uptake of ARGs by aiding the transfer of plasmid-mediated ARGs^58^ and zinc can increase ARG abundance by promoting HGT^59^. However, contrary to the negative correlation identified in this study, some studies found a significant positive correlation between potassium and the coexistence of soil multidrug resistance genes^60^. These conflicting findings suggest that the relationship between potassium and ARG richness may be bacterium-dependent. Another noteworthy environmental variable linked to ARG richness was soil pH. Studies have consistently demonstrated that soil pH is a crucial determinant affecting the diversity and composition of ARGs^61^. Our soil samples generally exhibited slightly acidic conditions (mean pH ± SD 6.6 ± 1.1), and we observed a negative correlation between pH and ARG richness, suggesting that lower soil pH levels are associated with higher ARG richness. This negative correlation might be attributed to the relationship between pH and HGT, where studies reported that acidic pH conditions promote the potential of HGT, while alkaline pH conditions attenuate HGT^62, 63^, consequently leading to a change in ARG richness. Given the complexity of natural environments and the co-correlations previously detected between soil property variables used in this study (e.g., between aluminum with calcium, magnesium, manganese, moisture, molybdenum, organic matter, total carbon, total nitrogen, and zinc)^29^, further experiments are needed to better understand the role of soil properties in influencing ARG richness.

Besides soil properties, surrounding land use, particularly forest coverage, was found to be positively associated with ARG richness in soil-dwelling *Listeria*. This is consistent with a previous global study which indicated that forests, irrespective of their location and type (boreal, cold, temperate, or tropical), exhibit the highest richness of ARGs in their soils^4^. The elevated presence of ARGs among *Listeria* in soil environments where surrounding areas have higher coverage of forest may be connected to the movement of wild animals, such as deer^64^, bird^65^, reptiles^66^, and rodents^67^ which serve as carriers of ARGs. These genes are subsequently excreted into the soil through fecal matter, contributing to the process of ARG acquisition.

Surrounding land use was also found to be a key driver of the diversification of ARG sequences in soil-dwelling *Listeria*. Prior research has highlighted the contribution of land use to the evolution of microbial antibiotic resistance^68^. For example, in cropland areas, the extensive use of antibiotics to enhance crop productivity^69^ selects specific ARGs. However, it is not clear how surrounding land use may affect the genetic diversification of ARGs. A plausible explanation is that surrounding land use could influence the soil properties of the natural environment, which indirectly impose selective pressures on the genetic diversification of ARGs. For instance, surrounding cropland and pasture coverage, in which we previously detected a positive correlation with soil magnesium^29^, were found to be associated with the sequence diversification of *mprF* and *norB* in this study. Moreover, we found evidence of PS in *mprF* and *sul*. There are four significant environmental variables exclusively associated with their sequence divergence, all of which are soil minerals (i.e., aluminum, copper, iron, and zinc). These minerals may induce oxidative stress by generating mineral-induced reactive oxygen species (ROS), which could stimulate genetic mutations^70^ and/or represent a selection pressure driving gene evolution^70^. Overall, the associations between surrounding land use and ARG richness and diversification detected in this study reflect potential anthropogenic effects on the dynamics of microbial antibiotic resistance in the natural environment.

In addition, we found the majority of environmental variables that were universally significantly and positively correlated with genetic divergence for all five ARGs in this study are climatic factors, including temperature and precipitation. It has been reported that increased temperatures can trigger adaptive responses in organisms, resulting in accelerated mutation rates, heightened genetic diversity, and enhanced genome plasticity^71^. Elevated temperatures directly contribute to higher mutation rates by promoting replication errors and inducing DNA damage^72^, thus elevating the mean strength of natural selection on genome-wide polymorphism^73^. Increased temperature also affects various biochemical and biophysical processes within cells^74^, leading to increased *de novo* genetic diversity. For instance, using enzyme kinetics theory, it has been shown that higher temperatures are likely to strengthen natural selection throughout the genome by amplifying the effects of DNA sequence variation on protein stability^73^. Alternatively, elevated temperatures might impact existing genetic variation by altering allele frequencies in genes or linked genomic regions associated with climate adaptation^75^.

## Conclusions

Through leveraging a national reconnaissance of *Listeria* isolates and further examination of the genetic context of their ARGs, we demonstrated the intrinsic nature of genetic antibiotic resistance traits predominately found in *Listeria sensu stricto* species in the soil environment. Considering the limited occurrence of prophages and plasmids carrying ARGs and the presence of a full set of functional competence genes in *Listeria sensu stricto* species, we propose that natural transformation may be the more plausible route for the HGT events observed in ARGs in soil-dwelling *Listeria*. In contrast, HGT of ARGs appears to be often achieved via conjugation in food and clinical isolates, suggesting that *Listeria* isolates from different environments may employ distinct HGT mechanisms for ARG acquisition. We also identified evidence of environmental selection likely triggered by soil properties, climate, and surrounding land use in the acquisition and diversification of ARGs in *Listeria*, highlighting the importance of monitoring the impact of environmental disturbance on the ecology and evolution of microbial antibiotic resistance. Overall, this study provides a baseline understanding of ecological and evolutionary processes governing ARG dynamics in soil environments and demonstrates *Listeria* as a model organism for elucidating the environmental factors that drive ARG mobilization and diversification across both *Listeria* pathogenic and non-pathogenic species.

## Materials and methods

### *Listeria* isolates and environmental data

The genomic data of 594 *Listeria* strains collected from minimally disturbed natural environments across the contiguous United States, which were described in a previous study examining the mechanism underlying bacterial pangenome evolution^29^, was further analyzed to consider their carriage of ARGs. These genomes represent 19 *Listeria* species, predominantly *L. monocytogenes* (*n* = 177), followed by *L. welshimeri* (*n* = 141), *L. seeligeri* (*n* = 98), and *L. booriae* (*n* = 90). Other *Listeria* genomes in our dataset included *L. innocua* (*n* = 33), *L. marthii* (*n* = 14), *L. cossartiae* (*n* = 11), *L. immobilis* (*n* = 9), *L. farberi* (*n* = 5), *L. grandensis* (*n* = 3), *L. ivanovii* (*n* = 2), and *L. rocourtiae* (*n* = 2). Single genomes were available for *L. swaminathanii, L. grayi, L. aquatica, L. weihenstephanensis, L. portnoyi*, and *L. newyorkensis*. Among the species included in this study, *L. monocytogenes, L. seeligeri, L. marthii, L. ivanovii, L. welshimeri, L. innocua, L. cossartiae, L. farberi, L. immobilis*, and *L. swaminathanii* are classified as *Listeria sensu stricto* species (491 genomes total), while others are *sensu lato* species (103 genomes total)^76^.

Previously reported environmental metadata encompassing 34 variables^29^ paired with this genomic dataset were also examined further in the present study. These environmental variables include 3 spatial (latitude, longitude, and elevation), 17 soil properties (moisture, total nitrogen, total carbon, pH, organic matter, aluminum, calcium, copper, iron, potassium, magnesium, manganese, molybdenum, sodium, phosphorus, sulfur, and zinc), 4 climatic (precipitation, wind speed, maximum and minimum temperatures), and 10 surrounding land-use (open water, barren, forest, shrubland, grassland, cropland, pasture, wetland, and developed open space categorized as > 20% and < 20% impervious cover)^29^ variables.

### Detection of ARGs, competence genes, flagellar genes, and mobile genetic elements (MGEs)

ARGs, competence genes, and flagellar genes were identified through BLASTN searches^77^, using an E-value of 0.01 and without restrictions on percent identity, against a reference database sourced from the BIGSdb-Lm platform^35^. BIGSdb-Lm is a curated bacterial isolate genome sequence database specializing in *L. monocytogenes*^35^. A total of 25 ARGs, 12 competence genes, and 31 flagellar genes were extracted from the platform (**Supplementary Table 5**)^35^. The gene with the highest bit-score was chosen for further analysis. Subsequently, the presence of premature stop codons and sequence coverage (%) was assessed for each detected gene. Genes were categorized as putatively functional if their sequence coverage exceeded 80% and no premature stop codon was detected; non-functional if its sequence coverage ranged between 30% - 80% or premature stop codon was detected; and absent if sequence coverage was < 30% or no hits were observed in the BLASTN searches^78^. Based on this categorization, we further simplified the classification as present but not necessarily functional (referred to as “present gene” hereafter) and putatively functional (referred to as “functional gene” hereafter).

To minimize the possibility of false negative predictions of ARGs in *sensu lato* species potentially arising from dissimilarities in ARG sequences to those of *L. monocytogenes* available in the BIGSdb-Lm database, we conducted cross-validation using two approaches. First, we obtained the reference genome of *L. rocourtiae* FSL F6-920 (accession number: AODK01) from the NCBI database and predicted its ARGs using our approach described above. This strain was selected because it belongs to the *Listeria sensu lato* group, and genotypic and phenotypic antibiotic resistance data for this strain are available and published in literature^79^. Through this analysis, we identified three ARGs: *sul*, *mprF*, and *fosX*. *fosX* appears to be functional, aligning with its known phenotypic fosfomycin resistance^79^, while *sul* and *mprF* were categorized as “not functional” due to the presence of a premature stop codon (**Supplementary Table 6**). However, this strain was found to be phenotypically resistant to sulfamethoxazole (conferred by *sul*)^79^, which was inconsistent with prediction according to the above criteria in this study. The functionality of *sul* might be attributed to its high coverage (94.6%) and the position of the stop codon in a later part of the gene (**Supplementary** Fig. 5). Secondly, we BLASTed our genomes against the Comprehensive Antibiotic Resistance Database (CARD)^37^, which includes a more diverse set of reference ARGs than BIGSdb-Lm. A consistent trend and no significant differences in prevalences for present ARGs (paired two-sided t-test *P* = 0.81) and functional ARGs (paired two-sided t-test *P* = 0.11) were observed between the predictions from BIGSdb-Lm and CARD (**Supplementary** Fig. 6). In addition, BLAST against CARD did not identify any *sul*, an ARG commonly found in *L. monocytogenes*^23, 35^. In contrast, it predicted *fosXCC,* an ARG with a similar function to *fosX,* and is commonly identified in *Campylobacter coli* rather than *Listeria* species^80^. From this, we conclude that our prediction of ARGs in *Listeria* species using BLAST against BIGSdb-Lm is robust, sensitive, and conservative.

To predict MGEs, including plasmids, insertion sequences (ISs), transposons, and prophages, we employed specialized programs, including PlasmidFinder2^81^ (0.6 cutoff) to identify plasmids; ISEScan^82^ and ISAbR_finder^83^ to detect ISs and ISs associated with ARGs; TnFinder^83^ to identify composite transposons; and PHASTER^84^ for prophage prediction. All programs were employed with default settings if not specified. Subsequently, a comparison was made to determine if the functional ARGs are present in (for plasmids and prophages) or near (for IS, within 2000 bp^83^) the predicted MGEs, based on their genomic coordinates. Positive findings were visualized using Gene Graphics^85^.

### Richness, diversity, and the spatial distribution of ARGs

The richness and Shannon-Wiener diversity index of present and functional ARGs were computed for each genome using the skbio library in Python 3.6.8. Since a strong positive correlation between ARG richness and diversity was observed (Spearman ρ = 1, *P* < 10e- 30; **Supplementary** Fig. 7), subsequent analyses only focused on ARG richness. To evaluate the association between the ARG richness of *Listeria* species and their genetic similarity to *L. monocytogenes*, we averaged the previously reported pairwise average nucleotide identity (ANI)^29^ for each genome based on their respective species compared with one *L. monocytogenes* genome and correlated it with the average richness of ARGs (both present and functional) using Spearman’s rank correlation analysis. The distribution of ARG richness for *Listeria* species was visualized using Mercator Projection and the Basemap Matplotlib Toolkit v.1.2.1 in Python v.3.6.8.

### Phylogenetic tree annotation, ARG tree construction, and PS detection

We annotated a previously published core single-nucleotide polymorphisms (SNPs)- based phylogenetic tree of 594 *Listeria* genomes^29^ with details about species, ARGs, plasmids, competence genes, flagellar genes, and the proportions of ISs, transposons, and prophages. The annotations were conducted using the Interactive Tree of Life (iTOL) webserver^86^.

To investigate the potential shared ancestry of ARGs within *Listeria*, we built a gene tree for each ARG. First, we aligned the nucleotide sequences of each ARG using MUSCLE v.3.8.31^87^. Then, we built the trees using a maximum likelihood method with 1,000 bootstraps in IQ-TREE^88^. IQ-TREE implements ModelFinder to select the best evolutionary model for the phylogenetic estimates^89^. The best-fit models, determined by the Bayesian information criterion (BIC), were TN+F+I+R4, GTR+F+I+R5, K3Pu+F+I+R3, HKY+F+I+R3, and TVM+F+I+R4 for *lin, mprF, sul, fosX*, and *norB*, respectively. All gene trees were then visualized through the iTOL webserver^86^.

To assess whether PS occurs across the entire gene in the functional ARGs, we utilized the BUSTED (branch-site unrestricted statistical test for episodic diversification) model in HyPhy^90^. A likelihood ratio test (LRT) was performed, comparing two models: the unconstrained model (allowing for PS) and the constrained model (disallowing PS). Statistical significance was determined by approximating the test statistic to a χ2 distribution. ARGs with a *P* < 0.05 were considered to exhibit evidence of PS, at least at one specific site in the ARG.

### Assessment of the relationships between environmental variables and the richness and diversification of ARGs

Two-sided Mann–Whitney *U* tests were employed to identify significant differences for each of the environmental variables between samples positive for *Listeria sensu stricto* and *sensu lato* species followed by a Benjamini-Hochberg (BH) false discovery rate (FDR) adjustment to account for multiple testing. An FDR-adjusted Spearman’s partial correlation analysis, controlling for genetic similarity, was performed to evaluate associations between ARG richness and each environmental variable. This approach addresses potential confounding effects of genetic similarity to *L. monocytogenes*. Environmental variables with an FDR-adjusted *P* < 0.05 were considered significant. Following this, a variation partitioning analysis (VPA) was conducted, controlling for genetic similarity to *L. monocytogenes*, to evaluate the relative contribution of each group with significant environmental variables (i.e., geolocation, soil property, climate, or land use) to ARG richness. Both genetic similarity and environmental data were structured in matrix format, enabling the use of adjusted *R*^2^ in redundancy analysis (RDA) ordination to partition the variation. VPA was executed and the respective adjusted *R^2^*value was visualized as a Venn diagram using the ’varpart’ function in the vegan package v.2.6-4 within the R environment. Furthermore, multidimensional scaling (MDS) along with a permutational multivariate analysis of variance (PERMANOVA) test was used to compare overall environmental conditions for isolates with and without functional ARGs. PERMANOVA *P* < 0.05 indicates that there are significant differences in the environmental conditions between groups of isolates with and without functional ARGs.

A series of Mantel tests were conducted to assess the relationships between the distance matrices of environmental variables and ARG sequences. Briefly, genetic dissimilarity between genomes for a given ARG was quantified using the Levenshtein distance^91^, while dissimilarity between genomes for a given environmental variable (excluding longitude and latitude)^29^ was calculated using Euclidean distance. Geographic distance was computed based on longitude and latitude using the haversine formula. FDR adjustment was applied to correct multiple testing. VPA was then performed using the calculated distance matrices to assess the relative contribution of each group with significant environmental variables identified to the sequence diversification of ARG.

### Machine learning models to predict the presence of ARGs using environmental variables

To predict the presence of ARGs with environmental variables as features, we developed an end-to-end machine learning-based framework that embodies a series of individual software programs (e.g., scikit-learn and SHapley Additive exPlanations, SHAP) written in Python for data preparation, hyperparameter tuning, model training, and testing, model evaluation through cross-validation, and visualization. Samples were first cleaned and split into the training set (80%) and the testing set (i.e., the holdout set; 20%) in a stratified fashion. The training set was further split into 5 stratified folds for cross-validation, in which a collection of predefined models (i.e., classifiers based on decision trees, random forest, multilayer perceptron, support vector machines, and gradient boosting) were trained and tested with a random set of hyperparameters (**Supplementary Table 4**). The average area under the receiver operating characteristic curve (auROC) score was used to evaluate the performance of the models across the 5 rounds of cross-validation. To account for stochasticity introduced by the random splitting of samples and division of training data into 5 folds, we repeated these steps 10 times. We selected the best model and its hyperparameter set with the highest interquartile mean of the auROC scores out of the 10 repetitions among the predefined models. The interquartile means of the auROC and area under the precision-recall curve (auPR) scores of the best model that was exclusively trained on the training set were reported based on a single evaluation of the holdout data from each of the 10 repetitions. The importance of the features was quantified using SHAP^43^.

## Data availability

The data used in this study, including *Listeria* genomic data and environmental metadata, were previously published in Liao et al, 2021^29^.

## Code availability

Code to replicate all analyses is available at https://github.com/leaph-lab/Soil_Listeria_ARG_manuscript.

## Acknowledgments

We thank all members of the Liao laboratory for their enriching discussions. This work was funded by the Virginia Tech Center for Emerging, Zoonotic, and Arthropod-borne Pathogens (CeZAP) Interdisciplinary Team-building Pilot Grant (JL) and the Infectious Disease Interdisciplinary Graduate Education Program (ID IGEP) fellowship (YG). The funder played no role in the study design, data collection, analysis and interpretation of data, or the writing of this manuscript.

## Author contributions

JL designed the study. YG, BA, AN, HZ, and JL analyzed the data. YG wrote the paper with input from JL, MP, LK, and AP.

## Competing interest

All authors declare no financial or non-financial competing interests.

## Supplementary Figures

**Supplementary Fig. 1.**
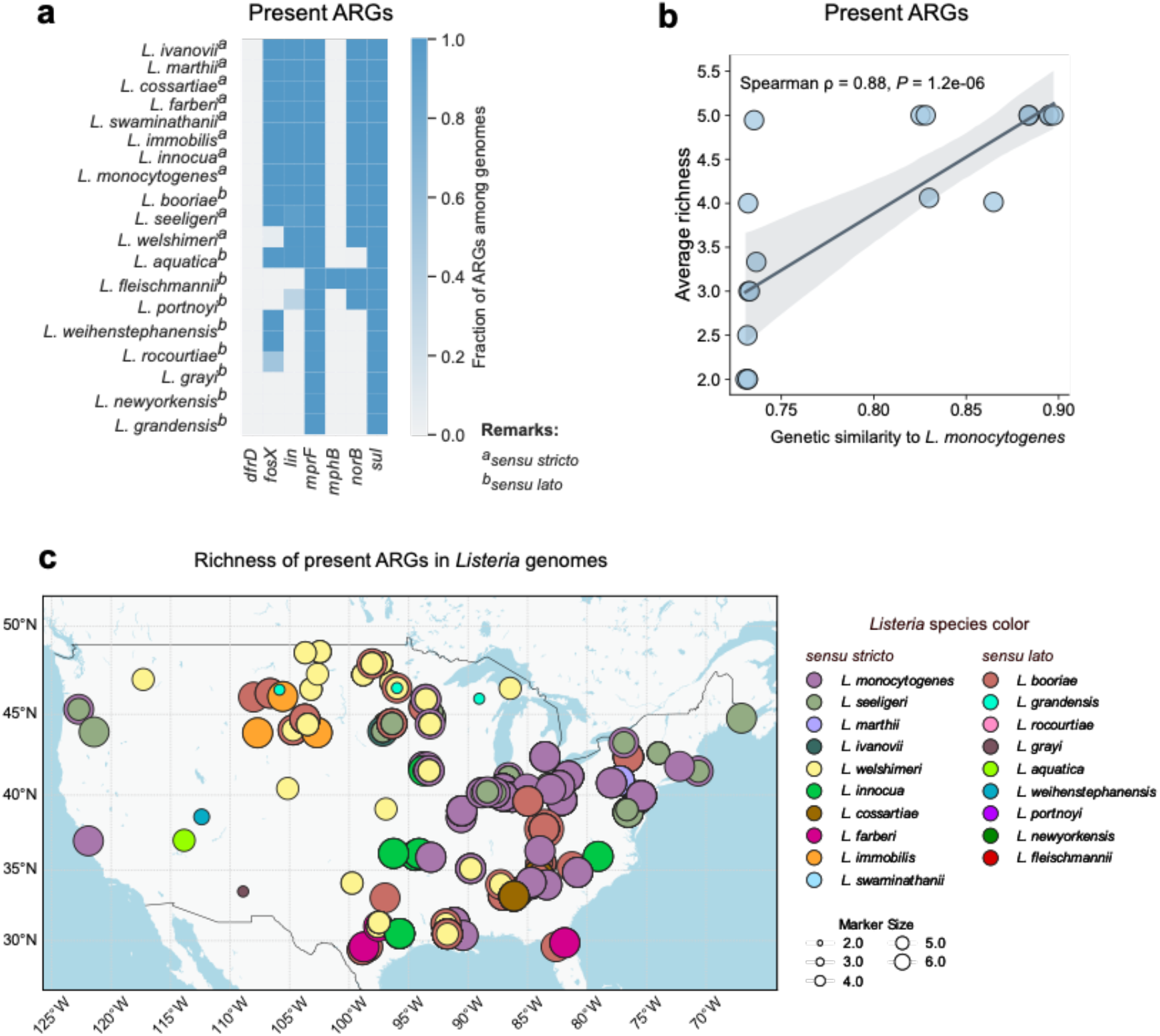
| ARG profiles among soil-dwelling *Listeria* and their national distribution. a. Propor4on of present ARGs among different *Listeria* species. **b** Correla4on between gene4c similarity to *L. monocytogenes* and average richness of present ARGs. Gene4c similarity was calculated based on pairwise ANI between different *Listeria* species and *L. monocytogenes*. ρ represents the Spearman’s rank correla4on coefficient. **c** Richness of present ARGs among *Listeria* genomes across the United States. Circles and crosses indicate genomes with and without ARGs, respec4vely, and are color-coded by species. Circle size is propor4onal to the ARG richness calculated in each genome.

**Supplementary Fig. 2.**
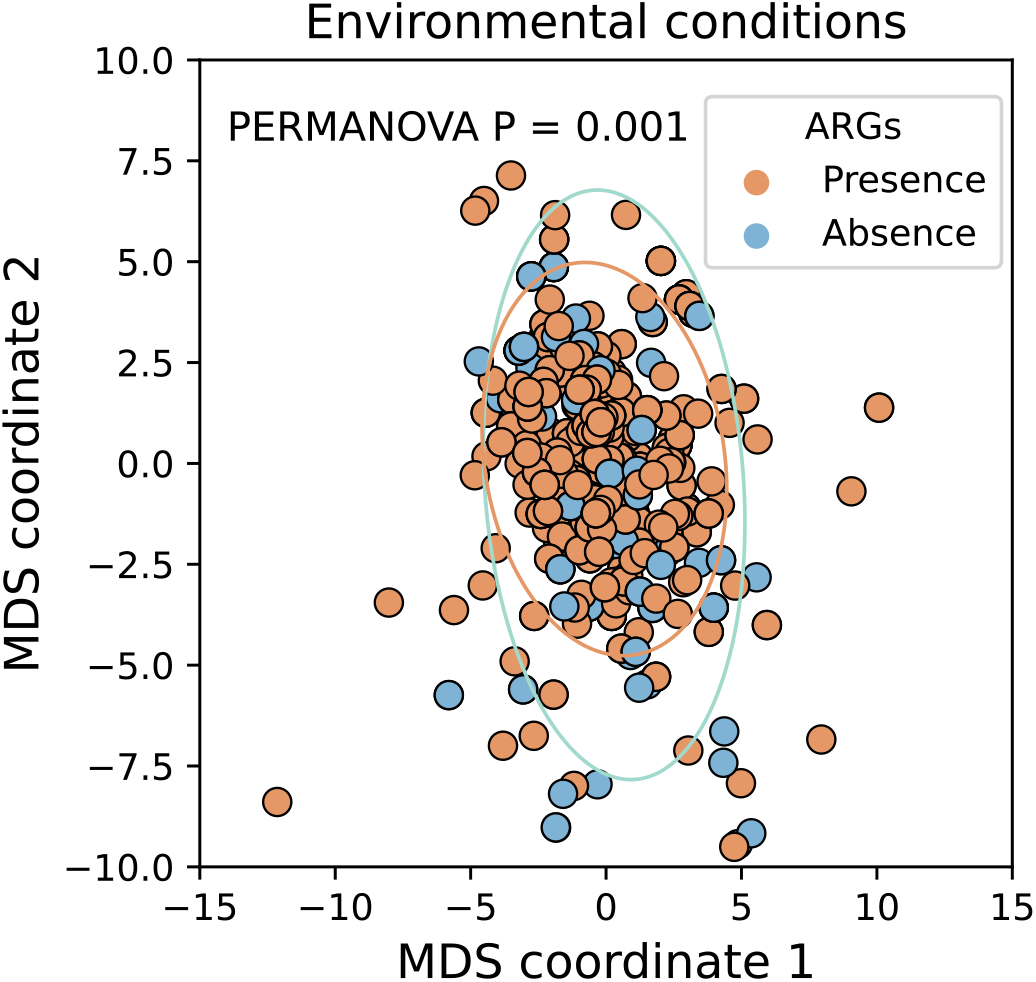
Mul4dimensional scaling (MDS) analysis for genomes with (red) and without (blue) ARGs based on environmental condi4ons. PERMONOVA *P* = 0.001 indicated that the environmental condi4ons were significantly different for genomes with and without ARGs. The size of the ellipse is determined by two 4mes the standard devia4on from the mean.

**Supplementary Fig. 3.**
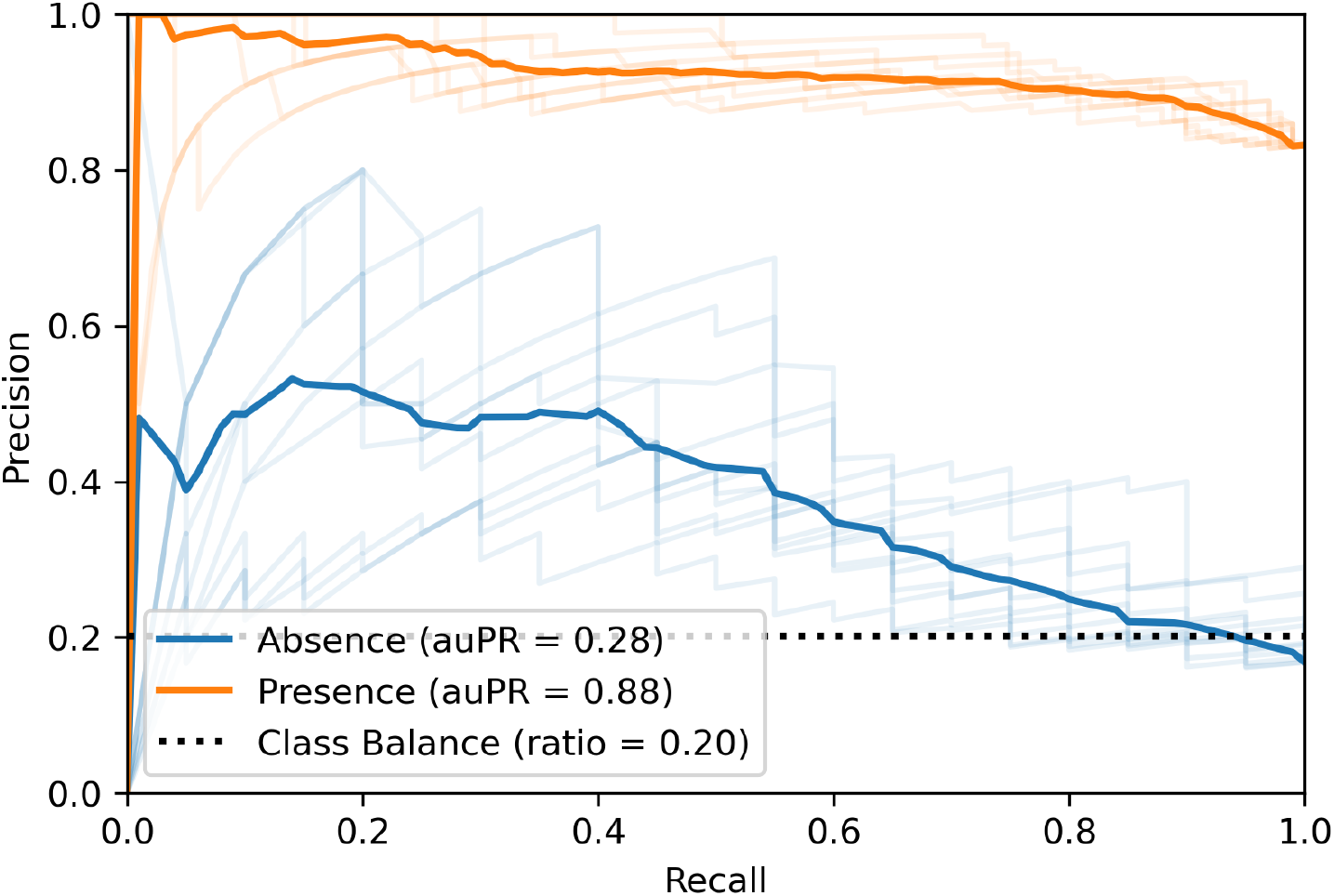
ARG predic4on precision of random forest model based on environmental variables (mean auPR = 0.88). The light-colored lines show PR curves for models trained only on the training set. Each curve reflects one evalua4on using holdout data, repeated 10 4mes. The dark-colored line represents the mean performance across these repe44ons. This model has high precision and recall for iden4fying ARG instances based on environmental data.

**Supplementary Fig. 4.**
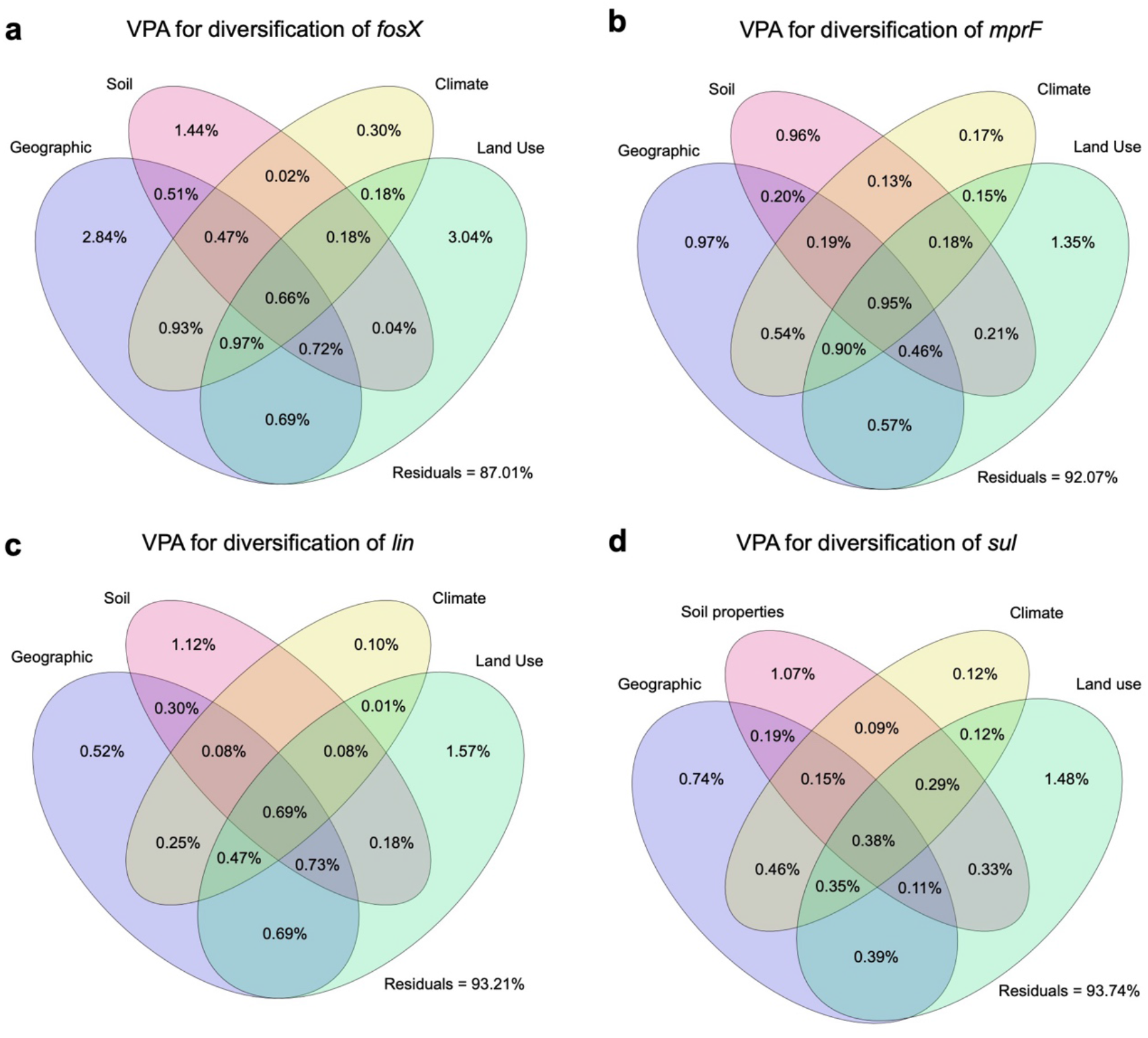
Venn diagram of varia4on par44oning analysis (VPA) showing the varia4on of the gene4c divergence of **a** *fosX*, **b** *mprF*, **c** *lin*, and **d** *sul* explained by environmental variable groups, including geographic loca4ons, soil proper4es, climate, and surrounding land use. Residuals indicate unexplained varia4on.

**Supplementary Fig. 5.**
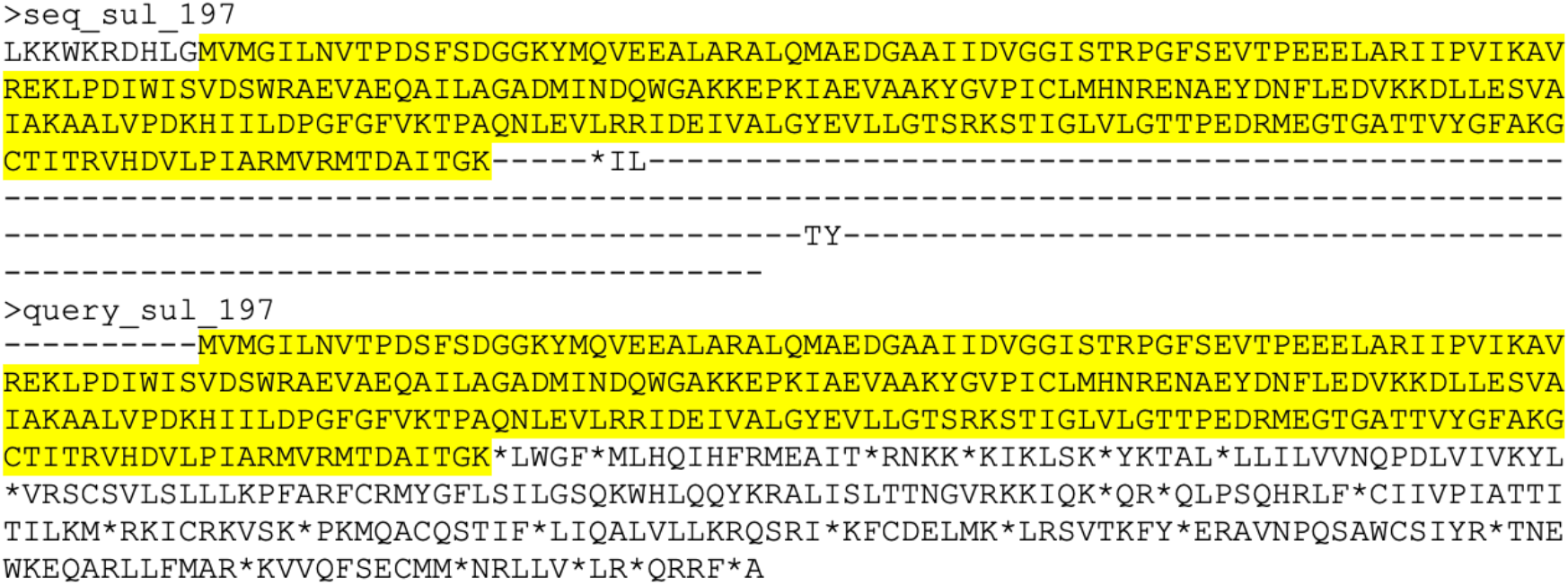
Amino acid sequence alignment of Sul. DNA sequences for *sul* were retrieved from the BIGSdb-Lm database (seq_sul_197) and the BLASTN output (query_sul_197), respectively. These sequences were then translated into amino acids and aligned using MEGA11 software^92^ to detect the position of the stop codons. The conserved region for amino acids was highlighted in yellow, and “*” indicates the identified stop codons.

**Supplementary Fig. 6.**
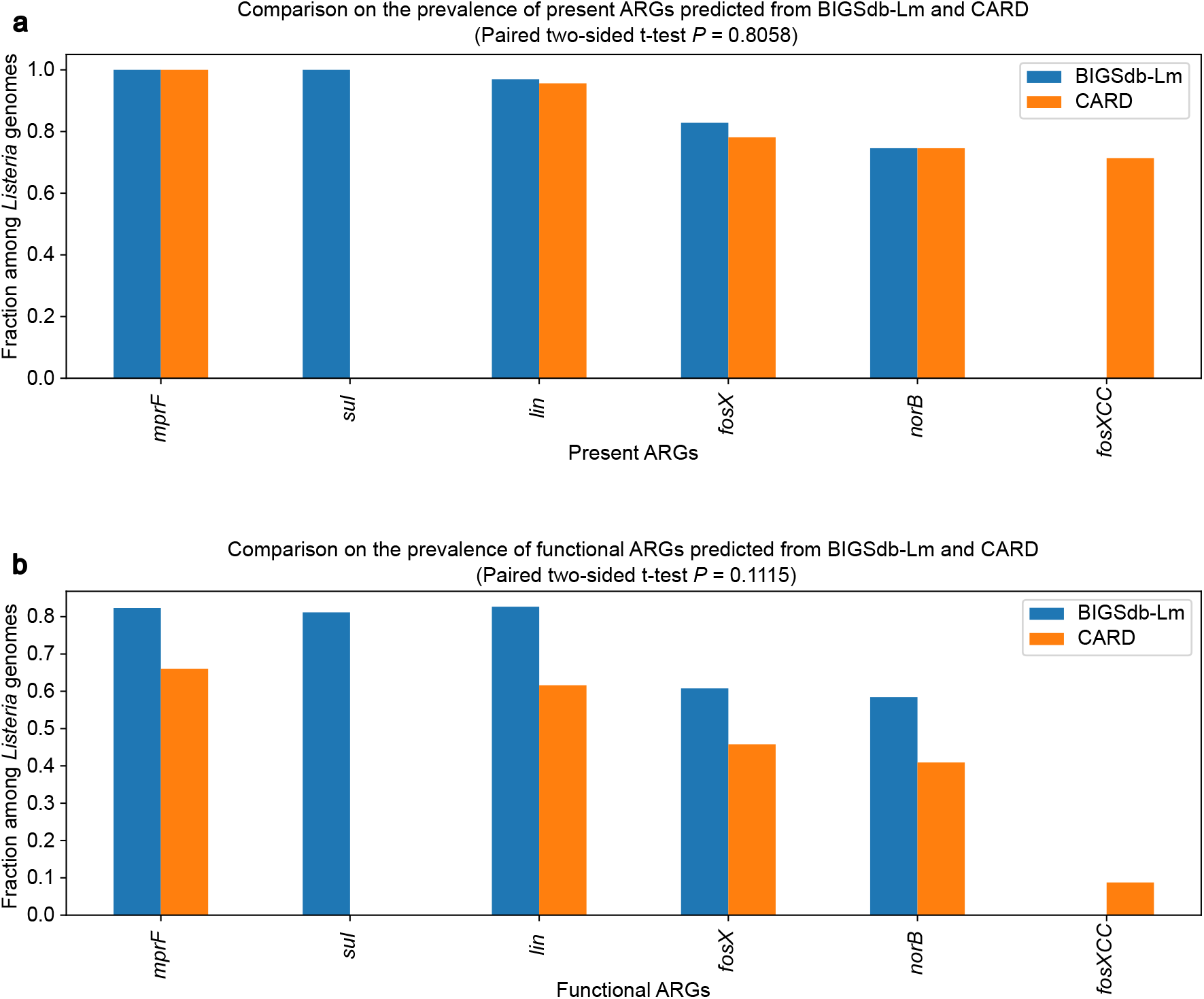
Comparing the prevalence of **a** present and **b** functional ARGs between the predicted outputs from BIGSdb-Lm and CARD using paired two-sided t-test revealed a consistent trend. No significant differences in prevalences were observed between the predictions from BIGSdb-Lm and CARD. BLASTN against CARD did not identify any *sul*, and BLAST against BIGSdb-Lm did not predict *fosXCC*. It is noteworthy that *fosXCC*, a gene resistant to fosfomycin with a similar function to *fosX*, was commonly identified in *Campylobacter coli* rather than *Listeria* species^80^.

**Supplementary Fig. 7.**
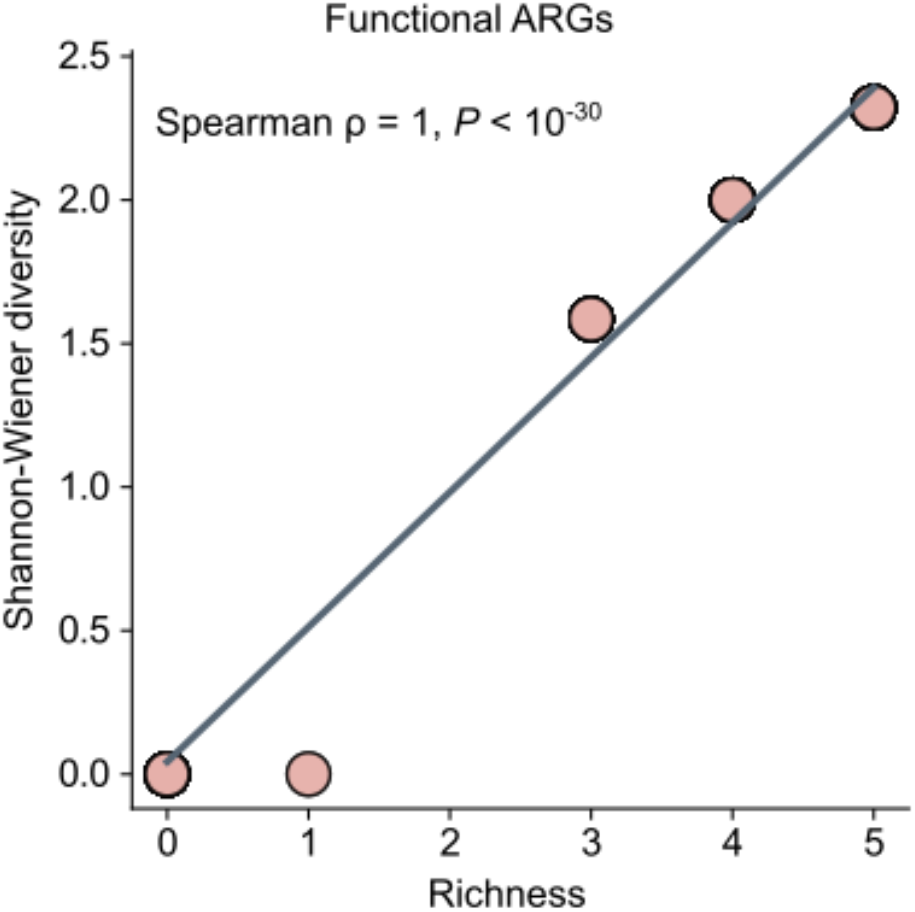
Spearman’s rank correlation between the richness and diversity of functional ARGs. ρ represents the Spearman’s rank correlation coefficient. This suggests a strong positive correlation between ARG richness and diversity.

## Supplementary Tables

**Supplementary Table 1** List of incomplete prophages identified by PHASTER, along with the associated ARGs they contain.

**Supplementary Table 2** List of plasmids predicted in the environmental *Listeria* isolates.

**Supplementary Table 3** Statistical comparison for the abundance of motility and competence genes between *Listeria sensu stricto* and *Listeria sensu lato* species.

**Supplementary Table 4** Hyperparameters for machine learning models.

**Supplementary Table 5** List of ARGs, motility genes, and competence genes retrieved from BIGSdb-Lm for BLASTN searches.

**Supplementary Table 6** BLASTN predicted output for *L. rocourtiae* FSL F6-920 (accession number: AODK01).

## Notes

### Competing Interest Statement

The authors have declared no competing interest.

